# Cognitive functions and underlying parameters of human brain physiology are associated with chronotype

**DOI:** 10.1101/2021.06.08.447543

**Authors:** Mohammad Ali Salehinejad, Miles Wischnewski, Elham Ghanavati, Mohsen Mosayebi-Samani, Min-Fang Kuo, Michael A. Nitsche

**Affiliations:** Department of Psychology and Neurosciences, Leibniz Research Centre for Working Environment and Human Factors, Dortmund, Germany; International Graduate School of Neuroscience, Ruhr-University Bochum, Bochum, Germany; Donders Institute for Brain, Cognition, and Behaviour, Radboud University Nijmegen, The Netherlands; Department of Psychology, Ruhr-University Bochum, Bochum, Germany; Institute of Biomedical Engineering and Informatics, Ilmenau University of Technology, Ilmenau, Germany; University Medical Hospital Bergmannsheil, Department of Neurology, Bochum, Germany

**Keywords:** Cognition, brain, chronotype, cortical excitability, neuroplasticity, non-invasive brain stimulation

## Abstract

Circadian rhythms have natural relative variations among humans known as chronotype. Chronotype or being a morning or evening person, has a specific physiological, behavioural, and also genetic manifestation. Whether and how chronotype modulates human brain physiology and cognition is, however, not well understood. Here we examine how cortical excitability, neuroplasticity, and cognition are associated with chronotype in early and late chronotype individuals. We monitor motor cortical excitability, brain stimulation-induced neuroplasticity, and examine motor learning and cognitive functions at circadian-preferred and non-preferred times of day in 32 individuals. Motor learning and cognitive performance (working memory, and attention) along with their electrophysiological components are significantly enhanced at the circadian-preferred, compared to the non-preferred time. This outperformance is associated with enhanced cortical excitability (prominent cortical facilitation, diminished cortical inhibition), and long-term potentiation/depression-like plasticity. Our data show convergent findings of chronotype can modulate human brain functions from basic physiological mechanisms to behaviour and higher cognitive functions.

## Introduction

Circadian rhythms are basic, daily cyclical processes that affect a wide range of physiological and behavioural manifestations and show significant variations in the human population^1^. Circadian *preference* or “chronotype”, describes an individual’s physical and behavioural preference for earlier or later sleep timing as a result of coupling between internal circadian cycles and the need for sleep^2^. The modulatory effects of circadian rhythms on basic physiological processes (e.g. cell cycle, body temperature, sleep-wake cycle) in living organisms are well established. Research performed during the last two decades has been primarily dedicated to molecular and cellular links between circadian rhythms and respective physiological processes in mammals, including humans^3, 4^. In recent years, respective research interest was broadened to fields such as genetics^2^, brain physiology^5^, and cognition^6, 7^.

This renewed interest in the “time-of-day” and “circadian rhythm” effects on human brain physiology and cognition is fueled by technological advances in human cognitive neuroscience^5,6^. Given that modern lifestyle is becoming less dependent on the 24-hr day-night cycle, an increased understanding of how the human brain and cognitive functions are influenced by chronotype and optimal time-of-day, has broad implications for human wellbeing, public health, working environments, school performance^8^, and disease-related pathophysiology^9-11^. Here we explored the interaction of chronotype and time-of-day effects on those aspects of human brain physiology, including cortical excitability and neuroplasticity, that determine adaptive behaviour in both healthy humans and clinical populations. We also investigated motor learning and higher-order cognitive functions, such as attention and working memory, and their associations with respective physiological processes, to reveal mechanisms of chronotype-dependent performance differences.

Technological advances in the neurosciences introduced non-invasive brain stimulation (NIBS) as a safe method for studying and directly modifying brain functions in humans^12^. Several NIBS techniques and protocols, including transcranial magnetic stimulation (TMS) and transcranial electrical stimulation (tES) are widely used to non-invasively *monitor* and *induce* changes of cortical excitability, and neuroplasticity^12, 13^ that underlie behaviour and cognition. Cortical excitability refers to responsiveness and response selectivity of cortical neurons to an input processed by the brain and is, therefore, a fundamental aspect of human brain functioning and cognition^5, 14^. TMS, which is based on principles of electromagnetic induction, can be applied in different paradigms to measure various aspects of cortical excitability^15^. These paradigms provide information about different neurotransmitter systems involved in cortico-cortical and corticospinal *excitability* (e.g. glutamatergic, dopaminergic, GABAergic, cholinergic systems). Monitoring cortical excitability with TMS enhances our understanding of the physiology of brain functions and cognition^12, 15^ as well as basic synaptic mechanisms involving long-term potentiation (LTP) or long-term depression (LTD)-like plasticity^13^.

Cortical excitability can be also modulated via induction of LTD/LTD-like plasticity, providing novel opportunities for examining a specific and mechanistic contribution of cortical regions to human behaviour^16, 17^. Transcranial direct current stimulation (tDCS) is a tES technique that can modulate and induce changes of cortical excitability via a weak, painless electrical current applied to the scalp^12, 18^. TDCS effects on cortical excitability are polarity-specific, with anodal stimulation inducing LTP-like plasticity and cathodal stimulation inducing LTD-like plasticity at the macroscopic level in humans^19, 20^. Mechanisms of plasticity induction via tDCS were demonstrated in previous animal^21, 22^ and human studies. These mechanisms are based on alterations of resting membrane potentials (for the acute effects) as well as glutamatergic, GABAergic, and calcium alterations, involving NMDA and AMPA receptors (for LTP LTD-like plasticity)^20, 23^. Both, LTP and LTD-like processes are assumed important physiological substrates of learning and memory formation^17^. In this line, tDCS has been shown to modulate learning and memory formation^24^. Accordingly, if the propensity to develop neuroplasticity in the brain is modulated by chronotype, we expect to see respective effects on behaviour, especially learning, and memory formation.

Animal studies show a strong circadian impact on hippocampal plasticity and LTP^25, 26^. Similarly, neural excitability in invertebrates^27^ and cortical excitability in the human motor cortex^28, 29^ are modulated by circadian rhythms. However, the impact of circadian *preference* on human cortical excitability and respective cognitive functions, and also brain plasticity and learning and memory formation are not well-studied. Increased understanding of respective mechanisms is important, not only for extending basic knowledge of human brain functions but also because of the broader implications and applications to our daily life circumstances, such as working and educational environments. In this study, we first systematically investigated the impact of chronotype and time-of-day effects on *cortical excitability* and stimulation-induced *neuroplasticity* in the model of the human motor cortex. In the next step, we explored how chronotype affects performance on a *motor learning* task which is associated with motor cortical plasticity^30^. Finally, we investigated potential effects of chronotype on higher-order *cognitive functions* that are dependent on cortical excitability and usually controlled by non-motor areas (e.g. prefrontal cortex). In all behavioural tasks, we recorded electroencephalography (EEG) to further explore electrophysiological correlates of cognition under different chronotypes and times of the day. All measurements were conducted on two groups of “early-chronotypes (ECs)” (i.e., morning-type), and “late-chronotypes (LCs)” (i.e., evening-types) at two fixed times in the morning and evening to capture circadian peaks and troughs at participants’ circadian preferred and non-preferred times (Fig. 1). The sleep/wake timing, amount of sleep, ambient light, and seasonal variations during the experiment were controlled for or taken into account (see Methods).

**Fig. 1.**
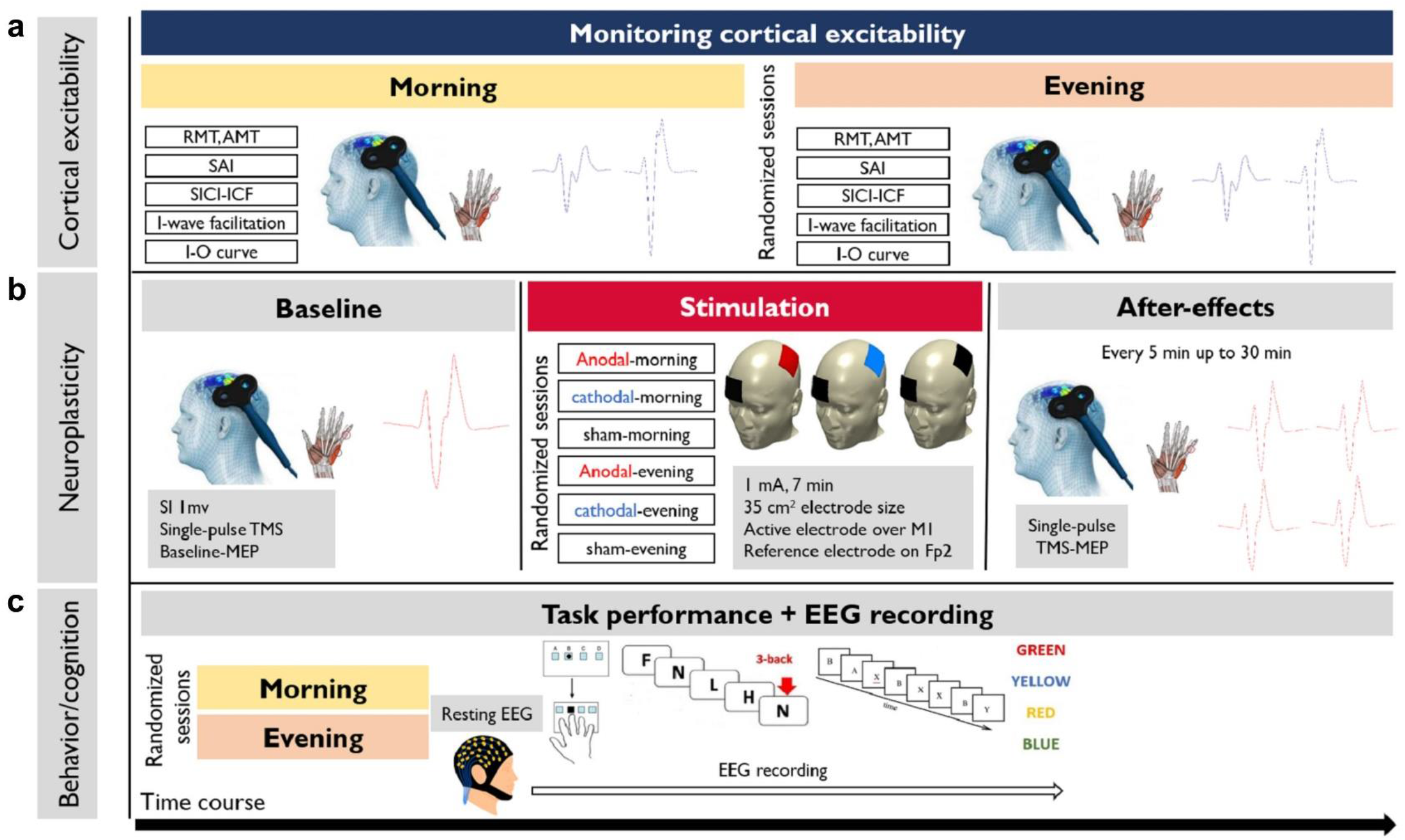
Course of study. **a** Cortical excitability was monitored once in the morning (8:30) and once in the evening (7:00 pm) at the same fixed time for all study sessions (one-week interval). Using single-pulse and double-pulse TMS protocols, cortico-spinal and cortico-cortical excitability was measured at the circadian-preferred and circadian non-preferred time. **b** each participant attended six sessions of tDCS (one session per week) in randomized order. TDCS sessions started at a fixed time in the morning and evening with a one-week interval between sessions. First, baseline cortical excitability was measured by inducing MEPs over the left M1 and measuring MEP of the target muscle (right ADM). Following a baseline measurement of 25 MEPs, 7 min of anodal, cathodal, or sham stimulation was delivered. MEP measurements were then conducted immediately in epochs of every 5 min up to 30 min after tDCS. **c** Following the resting-EEG acquisition, participants performed motor learning and cognitive (working memory, attention) tasks in two randomly-assigned sessions at the same time in the morning and evening while their EEG was recorded (one-week interval). The order of tasks was counterbalanced across participants. All tasks (SRTT, N-back, Stroop and AX-CPT) were presented on a computer screen (15.6″in. Samsung) in a soundproof electro-magnetic shielded room during EEG recording.

## Results

### Enhanced corticospinal excitability, and cortical facilitation, but reduced inhibition at the circadian-preferred time

We first monitored cortico-spinal and intracortical excitability of the human motor cortex at circadian-preferred and non-preferred times. Unless otherwise stated in this article, circadian-preferred time refers to morning and evening for ECs and LCs and circadian non-preferred time refers to evening and morning for EC and LCs respectively. We obtained Input-Output curve (I-O curve), as a measure of global cortico-spinal excitability, and intracortical facilitation (ICF) as a measure of cortical facilitation. Short-interval cortical inhibition (SICI), I-wave facilitation, and short-latency afferent inhibition (SAI) were applied as cortical inhibition protocols. These TMS protocols are based on different neurotransmitter systems related to cortical facilitation (glutamatergic) and inhibition (GABAergic, cholinergic)^31-33^ (see Methods). Age, gender, and BMI did not covariate with MEPs obtained from TMS protocols in the ANOVA analyses (Table 1).

#### Input-output curve (I/O curve)

The I-O curve is a global measure of *cortico-spinal* excitability^34^ obtained by eliciting motor-evoked potentials (MEPs) at a range of different TMS intensities (see Methods). The slope of the I-O curve reflects excitability of cortico-spinal neurons modulated by glutamatergic activity at higher TMS intensities^34, 35^. The ANOVA results showed significant interactions of chronotype**×**daytime**×**TMS intensity (*F*_1.29_ =15.79, *p*=0.001; *η*p^2^ =0.36) and chronotype**×**daytime (*F*_1_ =25.43, *p*=0.001; *η*p^2^ =0.48) but no main effects of chronotype and daytime (morning vs evening) alone on the slope of the I-O curve (Table 1). Bonferroni-corrected post *hoc* comparisons showed that MEP amplitudes were significantly larger at 130% and 150% of resting motor threshold (RMT) intensity in the morning for ECs and in the evening for LCs compared to their circadian non-preferred time and the same time-point in the other group. Furthermore, MEP amplitudes at 110% of RMT intensity significantly increased only in the morning for ECs (Fig. 2a).

**Fig. 2.**
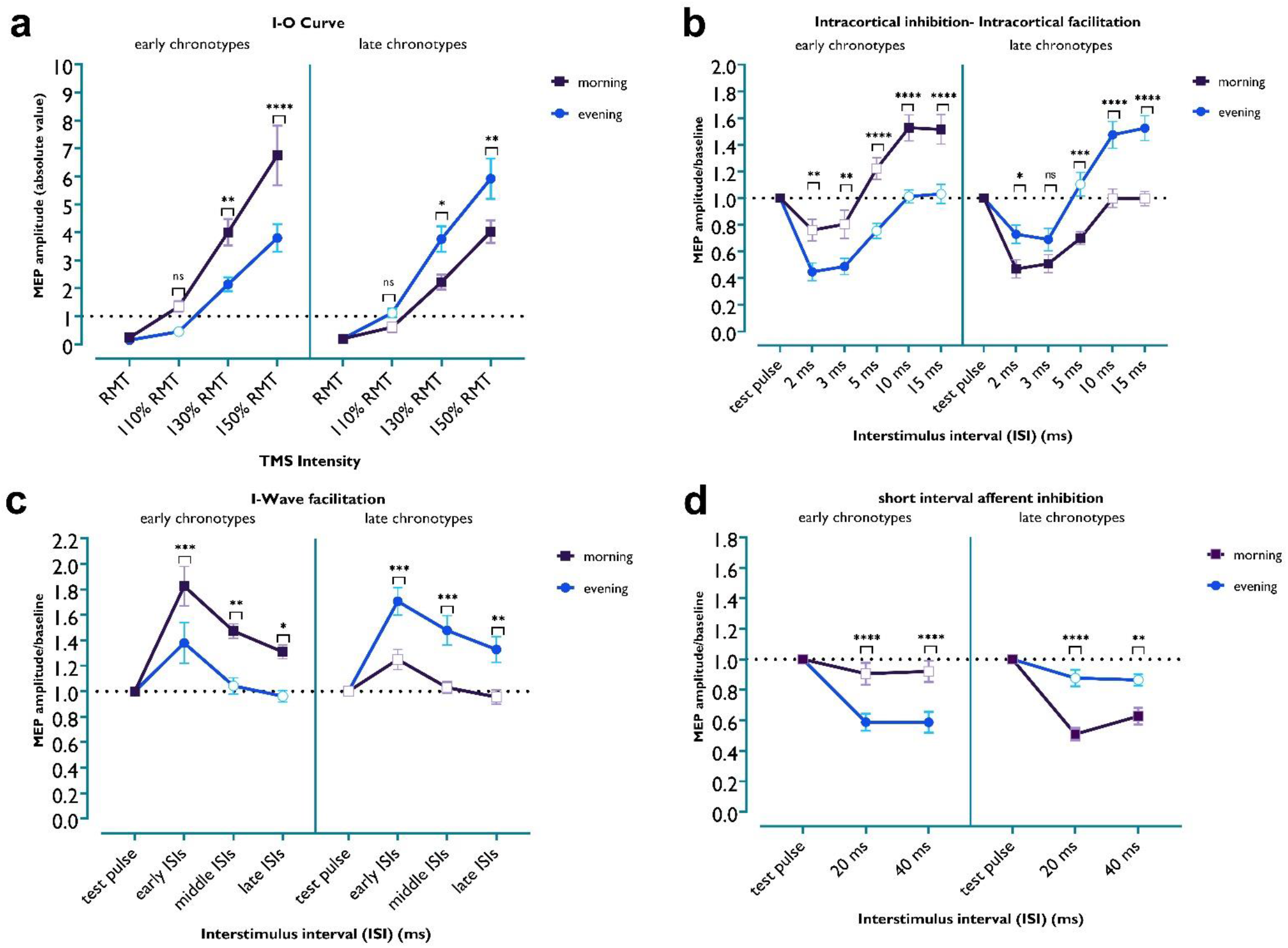
Monitoring cortico-spinal and corticocortical excitability with TMS protocols. **a** Global cortico-spinal excitability monitored by MEP amplitudes at different TMS intensities via the I-O curve protocol. ECs showed significantly higher cortico-spinal excitability in the morning than in the evening (*t*_*130%*_ =3.19, *p*=0.004; *t*_*150%*_ =5.054, *p*<0.001), and compared to the same time in LCs (*t*_*130%*_ =3.05, *p*=0.007; *t*_*150%*_ =4.67, *p*<0.001), and LCs display enhanced excitability in the evening (*t*_*130%*_ =2.64, *p*=0.026; *t*_*150%*_ =3.248, *p*=0.004), and compared to the same time in ECs (*t*_*130%*_ =2.78, *p*=0.017; *t*_*150%*_ =3.63, *p*=0.004) at higher TMS intensities. **b** Intracortical inhibition and facilitation measured by the SICI-ICF paired-pulse (pp)TMS protocol. Significantly higher cortical inhibition in the evening and morning were observed for ECs (*t*_*ISI2*_ =3.09, *p*=0.006; *t*_*ISI3*_ =3.13, *p*=0.005) and LCs (*t*_*ISI2*_ =2.57, *p*=0.031; *t*_*ISI3*_ =1.80, *p*=0.217), respectively. In contrast, cortical facilitation was significantly enhanced in the morning, and evening for ECs (*t*_*ISI10*_ =5.09, *p*<0.001; *t*_*ISI15*_ =4.79, *p*<0.001) and LCs (*t*_*ISI10*_ =4.71, *p*<0.001; *t*_*ISI15*_ =5.23, *p*<0.001), respectively. **c** I-Wave facilitation for monitoring GABA-dependent intracortical inhibition. Cortical excitability was significantly facilitated for early, middle and late ISIs in the morning for ECs (*t*_*early*_ =3.84, *p*=0.009; *t*_*middle*_ =3.70, *p*=0.001; *t*_*late*_ =2.992, *p*=0.018) and in the evening for LCs (*t*_*early*_ =3.92, *p*=0.007; *t*_*middle*_ =3.85, *p*=0.009; *t*_*late*_ =3.214, *p*=0.009), indicative for less cortical inhibition. **d** Inhibitory effect of peripheral nerve stimulation on motor cortical inhibition, as measured by SAI. The ECs showed more prominent cortical inhibition in the evening (*t*_*ISI20*_ =4.76, *p*<0.001; *t*_*ISI40*_ =4.99, *p*<0.001), whereas LCs showed more cortical inhibition in the morning (*t*_*ISI20*_ =5.50, *p*<0.001; *t*_*ISI40*_ =3.56, *p*<0.001). All pairwise comparisons were calculated using the Bonferroni correction for multiple comparisons. *n*=32 (16 per group). All error bars represent the standard error of means (s.e.m) across participants. Asterisks represent statistically significant comparisons.

*SICI-ICF*. In this double pulse TMS protocol, the interstimulus interval (ISI) between a subthreshold conditioning stimulus and a suprathreshold test stimulus determines inhibitory (ISIs 2 and 3 ms) or facilitatory (ISIs 10 and 15 ms) effects on cortical excitability^36^, which are reflected by a reduction or enhancement of MEP amplitudes (see Methods). The results of the ANOVA showed significant interactions of chronotype**×**daytime (*F*_1_ =72.16, *p*=0.001, *η*p^2^ =0.72) and chronotype**×**daytime**×**ISI (*F*_3.49_ =13.44, *p*=0.001, *η*p^2^ =0.33), but no significant effect of chronotype and time of day alone (Table 1). Bonferroni-corrected post *hoc* comparisons of the single pulse MEP revealed that both, ECs and LCs showed a significant increase of intracortical inhibition at ISIs of 2 and 3 ms at their circadian *non*-preferred time, compared with the single pulse-elicited MEP amplitudes (baseline) and respective ISIs at their circadian-preferred time (Fig 2b). At the circadian-preferred time, intracortical inhibition was significant only in the LCs. Regarding intracortical facilitation, MEP amplitudes at ISIs of 10 and 15 ms were significantly increased only at their circadian-preferred time in both groups, when compared with single pulse-elicited MEP amplitudes (baseline), respective ISIs at their circadian non-preferred time and the same time-point in the other group (Fig. 2b). Together, these results demonstrate a significantly *lower* cortical inhibition and *higher* cortical facilitation at the circadian-preferred time in both groups.

#### I-wave facilitation

Another method to monitor cortical inhibition is to explore facilitatory interaction between I-waves in the motor cortex that originate from corticospinal neurons^37^. In this TMS protocol, a suprathreshold stimulus is followed by a subthreshold second stimulus at different ISIs. I-wave peaks are mainly observed at three ISIs occurring at about 1.1-1.5, 2.3-2.9, and 4.1-4.4 ms after test pulse application^37^. We grouped these ISIs in epochs of early, middle and late ISIs and analyzed the MEP amplitude means. The overall ANOVA shows significant interactions of chronotype**×**daytime (*F*_1_ =44.24, *p*=0.001; *η*p^2^ =0.62) but no main effects of chronotype and time of day on I-wave peaks (Table 1). Bonferroni-corrected post *hoc* comparisons showed a significant increase of I-wave peaks for early, middle, and late ISIs, as compared to single-pulse MEPs in both groups at their circadian-preferred time of day. Moreover, I-wave peaks were significantly facilitated at the circadian-preferred time compared to the non-preferred time in each group and the same time in the other group (Fig. 2c). These results indicate an impact of chronotype on I-wave facilitation, indicative of reduced GABAergic inhibition at the circadian-preferred time.

*SAI*. SAI is a measure of cortical inhibition and reflects inhibitory modulation of the motor cortex via somatosensory inhibitory afferents. In this protocol, the TMS stimulus is coupled with peripheral nerve stimulation that has an inhibitory effect on motor cortex excitability at ISIs of 20 and 40 ms^38^. This inhibitory effect is linked to cholinergic^31^ and GABAergic^38^ systems. ANOVA results showed significant interactions of chronotype**×**daytime (*F*_1_ =114.20, *p* = 0.001; *η*p^2^ =0.62) and chronotype**×**daytime**×**ISI (*F*_1.61_ =30.10, *p*= 0.001; *η*p^2^ =0.52), but no significant main effects of chronotype and time of day (Table 1). Bonferroni-corrected post *hoc* comparisons revealed a significantly pronounced inhibitory effect of peripheral stimulation on cortical excitability at the circadian *non*-preferred time in both groups compared to the single TMS pulse. Moreover, cortical inhibition was significantly reduced in each group at their circadian-preferred time and between groups at the respective time points (Fig. 2d). This result is consistent with that of SICI, suggesting a reduction of cortical inhibition at circadian-preferred times.

Taken together, we monitored cortical excitability in ECs and LCs and found a strong dependence of motor cortical excitability parameters on chronotype and time-of-day, indicative for a prominent impact of these factors on neurotransmitter systems. When participants were at their circadian-preferred time, they showed higher levels of corticospinal excitability and cortical facilitation, and a lower level of cortical inhibition (Fig. 2), in accordance with a higher glutamatergic and lower GABAergic activity during the circadian-preferred, as compared to the non-preferred time of day. Neither the baseline measurements of protocols nor the stimulation intensity required to evoke MEP did differ across groups and times of day (Supplementary Tables 1,2). The results thus cannot be explained by different stimulation intensities across times of day.

### LTP/LTD-like plasticity in the motor cortex is facilitated at the circadian-preferred time in early and late chronotypes

Having demonstrated that cortical excitability in the motor cortex is chronotype-dependent, we were next interested in determining how the time-of-day dependent variation of cortical excitability affects LTP/LTD-like plasticity in early and late chronotypes. We predicted that motor cortical plasticity should be facilitated at the circadian-preferred time too. To test this hypothesis, we stimulated the primary motor cortex with anodal, cathodal and sham (control condition) tDCS (1 mA, 7 min, Fig. 3a) in each group at the same time in the morning and evening (6 sessions, weekly) and monitored neuroplastic effects of tDCS via single-pulse TMS with a fixed medium intensity before and after the intervention (see Methods). Depending on the stimulation polarity, tDCS results in LTP-like or LTD-like plasticity. With the chosen protocol, anodal tDCS enhances, while cathodal tDCS diminishes motor cortex excitability^39^ which can last for an hour or longer after tDCS^39, 40^. Analysis of blinding efficacy showed that participants could not discern between active and respective sham tDCS conditions (Supplementary Table 6). Side effects were minor and did not differ between intervention conditions except for the tingling and burning sensations (Supplementary Tables 4,5). Age, gender, and BMI did not covariate with TMS-induced MEP in the ANOVA analyses. ANOVA results showed significant interactions of chronotype×daytime stimulation×chronotype×daytime and stimulation×chronotype×daytime×timepoint (*F*_6.78_ =10.82, *p*=0.001; *η*p^2^ =0.28) but no interaction of stimulation×chronotype and timepoint×chronotype. The main effects of chronotype and time of day were not significant (Table 2).

**Fig. 3.**
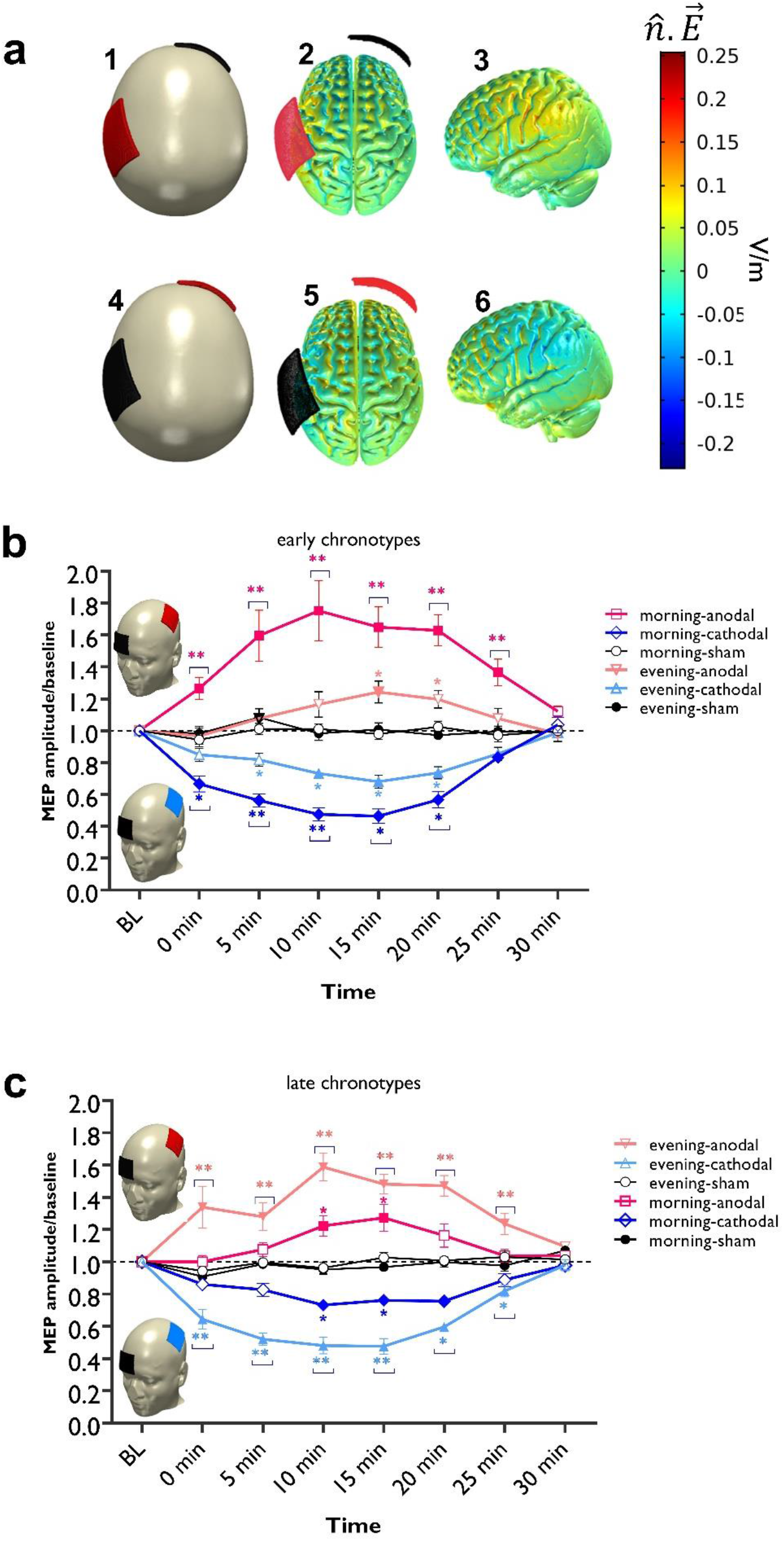
LTP/LTD-like plasticity induction in the motor cortex. **a** 3D model of the current flow distribution inside the head was calculated based on the finite element method using COMSOL Multiphysics software version 5.2 (COMSOL Inc., MA, USA). The electrical current flow induced by 1.0 mA stimulation intensity, and electrode positions C3-Fp2, for anodal (a1, 2,3), and cathodal (a4,5,6) stimulation over the motor cortex is shown. *ñ*.E refers to the absolute electrical field. **b, c** post-tDCS cortical excitability alterations after anodal, cathodal, and sham stimulation at the circadian-preferred and non-preferred times in ECs (b) and LCs (c) (*n* = 32). The results of the repeated-measures ANOVA showed significant interactions of stimulation**×**chronotype**×**daytime and stimulation**×**chronotype**×**daytime**×**timeline (Table 2). The main effects of time of day and chronotype were not significant, however they significantly interacted. Stimulation and timepoint did not significantly interact with chronotype or time of day (Table 2). Post *hoc* comparisons (Bonferroni-corrected) of MEP amplitudes to respective baseline values, the sham condition and respective stimulation conditions at different time of day are marked by symbols in the figures. Filled symbols indicate a significant difference of cortical excitability against the respective baseline values. The floating symbol [*] indicates a significant difference between the real vs sham tDCS conditions, and the floating symbol [**] indicates a significant difference between respective timepoints of tDCS condition at the circadian-preferred vs circadian non-preferred times. Sham stimulation did not induce any significant change of cortical excitability. All error bars represent the s.e.m. across participants.

#### Anodal stimulation

We analyzed the effects of anodal tDCS on MEP amplitudes compared to the baseline, across daytime and against sham condition via Bonferroni-corrected post *hoc t*-tests. For ECs, MEP amplitudes significantly increased at 5, 10, 15, 20 and 25 min after intervention in the *morning*, but only at 15 min in the evening, as compared to baseline MEP. Importantly, the increase of MEP amplitudes in the morning was significantly higher at all timepoints except for 30 min after the stimulation, when compared to the evening session and against the sham intervention (Fig. 3b). A reversed pattern of response was found for the LCs. Anodal tDCS significantly increased MEP amplitudes at 5, 10, 15, 20 and 25 min after stimulation in the *evening* and only at 10 and 15 min in the morning. The increase of MEP amplitudes in the *evening* was significantly higher when compared to the morning session and against the sham intervention for all time points except 30 min after stimulation (Fig. 3b).

#### Cathodal stimulation

Here, respective post *hoc* t-tests showed that cathodal tDCS resulted in a significant decrease of MEP amplitudes in both chronotypes at their circadian-preferred time compared to the baseline MEP and against sham at all times points except for 25 and 30 min after stimulation (Fig. 3c). Both groups showed a significant decrease of MEP amplitudes at 10, 15, and 20 min after cathodal stimulation at their circadian non-preferred time as well. However, the MEP decrease after stimulation was significantly larger in the morning for ECs (at 5 and 10 min) and in the evening for the LCs (0, 5, 10, 15), when compared to MEP size at the respective circadian non-preferred time (Fig. 3c).

Together, these results indicate that tDCS-induced LTP- and LTD-like plasticity of the motor cortex (after anodal and cathodal stimulation), which are dependent on glutamate and GABA activity^41^, were significantly stronger and longer-lasting in both, ECs and LCs at their circadian-preferred time. This aligns with our findings of higher cortical facilitation and lower cortical inhibition in the motor cortex at the circadian-preferred time, as described earlier.

#### Impact of chronotype on behavioural and electrophysiological aspects of motor learning

We found daytime-specific effects of chronotype on basic physiological functions of the motor cortex. LTP/LTD are important physiological foundations for learning and memory formation. Concentration of GABA^42^ and glutamate^17^ is important for motor learning and synaptic strengthening as well. If circadian preference modulates LTP/LTD processes and respective neurotransmitter systems, as shown in the previous section, better learning is expected at the circadian-preferred time. To this end, we investigated sequence motor learning using the serial motor learning task (SRTT) and its electrophysiological correlates and predicted enhanced motor learning at the circadian-preferred time. To test this hypothesis, participants in both groups performed SRTT at the same time in the morning and evening during EEG recording (see Methods).

We analyzed the differences in the standardized reaction time (RT) of block 5 vs 6, which is indicative of motor sequence learning acquisition, and block 6 vs 7 which is indicative of motor sequence learning retention (for analysis of absolute RT see supplementary Fig. 1). The 3×2×2 ANOVA results showed a significant interaction of block×chronotype×daytime (*F*_1.89_ =9.97, *p*<0.001; *η*p^2^ =0.27) but no interaction of block×chronotype, block×daytime, chronotype×daytime, or main effect of chronotype and daytime (Fig. 4, Table 3). Post *hoc* comparisons t-tests revealed that both groups significantly displayed longer RT at block 6 compared to block 5, indicative of sequence motor learning, at their circadian-preferred time (Fig. 4a,b). Importantly, the block 6-5 RT difference was significantly larger in both groups at their circadian-preferred time compared to the respective non-preferred time. To test if the learning sequence was preserved after the presentation of random stimuli in block 6, we analyzed RT differences of block 6 vs 7 too. The results showed that sequence learning was significantly retained at the circadian-preferred time as well (Fig. 4a,b and Supplementary material). Baseline block and block 6 RT, which contain stimuli in random order, did not significantly differ across morning and evening sessions in both groups (*F*_1_ = 3.39, *p*=0.076) and therefore, a generally slower RT at the circadian non-preferred time cannot explain RT differences in the learning blocks. Additionally, we analyzed the number of errors and RT variability in the learning blocks and found that both groups committed more errors at block 6 at their circadian non-preferred time (Supplementary Fig. S2a,b).

**Fig. 4.**
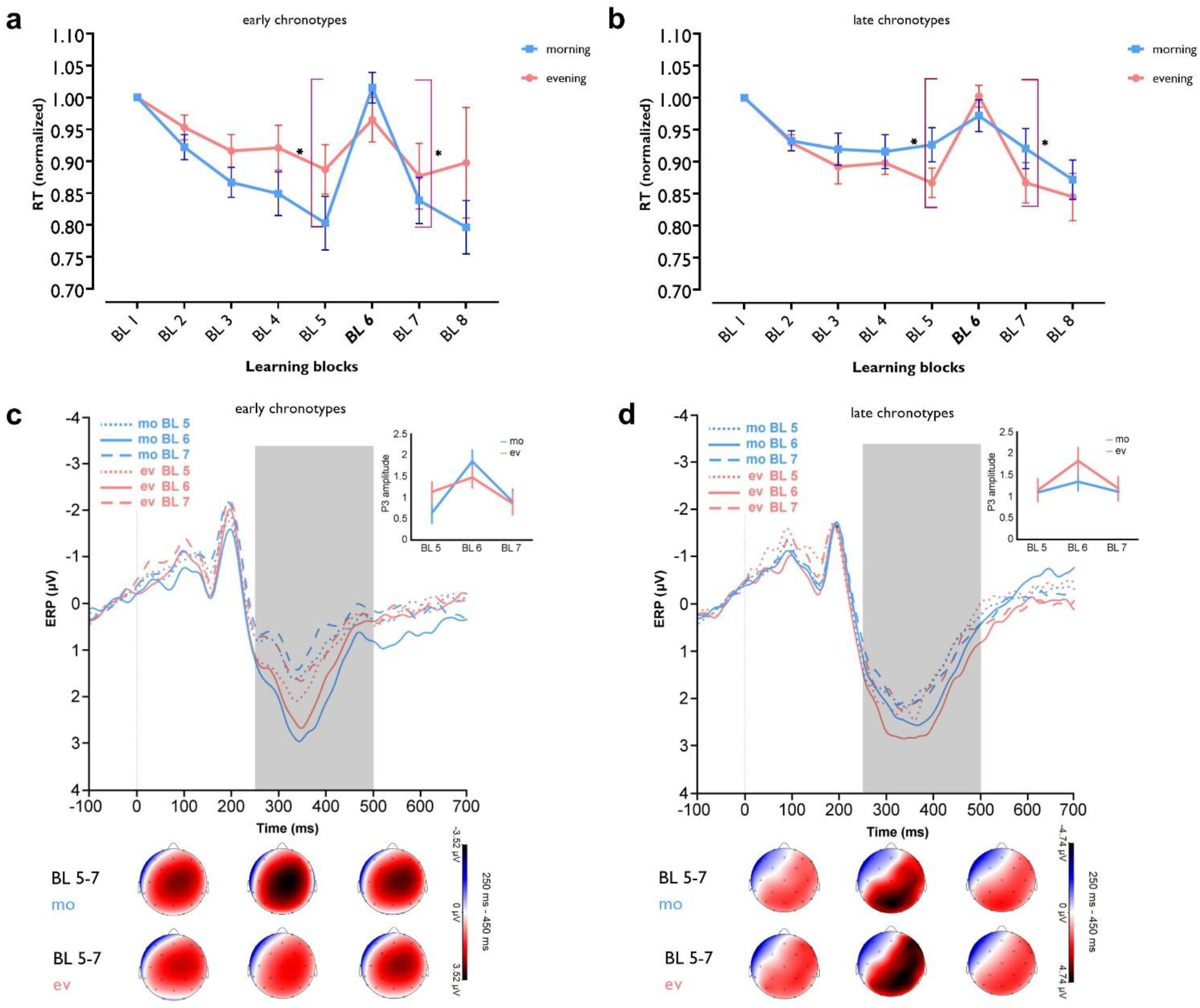
Chronotype affects motor learning performance and ERP correlates. **a, b**, The RT difference between blocks 5 and 6 mostly exclusively represents sequence learning. In ECs, the RT difference between these blocks was significant both in the morning and evening (*t*_*morning*_ =5.70,*p*<0.001, *t*_*evening*_ =2.93, *p*=0.010), but block 6-5 RT difference in the morning vs evening was significantly larger (*t*=2.90, *p*=0.012). In LCs, the respective RT difference was significant both in the evening and morning (*t*_*evening*_ =8.78, *p*<0.001; *t*_*morning*_ =2.40, *p*=0.029) and blocks RT difference was significantly faster in the evening, compared to the morning (*t*=2.72, *p*=0.016). The RT difference between blocks 6 and 7 was significant in the morning and evening for both ECs (*t*_*morning*_ =5.53, *p*<0.001; *t*_*evening*_ =3.39, *p*=0.004) and LCs (*t*_*evening*_ =5.06, *p*<0.001; *t*_*morning*_ =2.43, *p*<0.028). Blocks 6-7 RT difference in the morning vs evening was only marginally significant for LC (*t*=2.11, *p*=0.052). Asterisks [*] represent statistically significant differences between learning blocks RT (BL 6-5, BL 6-7]. The brackets refer to RT difference between blocks 6 vs 5 and 6 vs 7 in the morning for ECs and evening for LCs. **c, d** The P300 component of electrode Pz was calculated per block in both groups. Pairwise comparisons show that ECs displayed a significantly larger P300 at block 6 vs block 5 in the morning (*t*=4.63, *p*<0.001) vs evening (*t*=1.29, *p*=0.198), whereas LCs showed reversed results in the evening (*t*=3.22, *p*=0.001) vs morning (*t*=1.18, *p*=0.239). The P300 positivity of electrode Pz was significantly reduced at block 7 vs block 6 in ECs (*t*_*morning*_ =3.62, *p*<0.001; *t*_*evening*_ =2.33, *p*=0.022) and LCs (*t*_*morning*_ =1.11, *p*=0.269; *t*_*evening*_ =3.07, *p*=0.002) at their circadian-preferred time. All pairwise comparisons are calculated using *Student’s* t-test. *n*=31 (15 ECs, 16 LCs; data of one EC participant excluded due to sequence awareness). All error bars represent the s.e.m across participants. BL = block; mo = morning; ev = evening.

Next, we explored electrophysiological correlates of motor learning by analyzing event-related potentials (ERPs) of the learning blocks (see methods). The P300 component is evoked in response to stimuli of low probability and consists of the P3a (250-280 ms, reflecting frontal activity) and P3b (250-500 ms, reflecting temporo-parietal activity) waves^43^. It is affected by stimulus characteristics, including stimulus sequence^44, 45^ and is related to motor decision mechanisms^46^. We expected a higher-amplitude P300 component, when the learned sequence of stimuli is violated (random block 6), at the circadian-preferred times which resulted in superior motor learning. To test for statistical significance, we analyzed the P-300 amplitudes (250-500 ms) of blocks (5-7) and amplitude differences at block 5 vs 6 (sequence acquisition) and block 6 vs 7 (sequence retention). The ANOVA results revealed a significant interaction of block×chronotype×daytime (*F*_1.99_ =6.58, *p*=0.004; *η*p^2^ =0.18) for P-300 at the Pz electrode, but no significant main effects of chronotype and daytime (Fig. 4, Table 3). Post *hoc* comparisons confirmed our prediction and both, early and late chronotypes had a significantly larger P-300 amplitude in block 6 compared to blocks 5 and 7 (indicative of sequence learning) at their circadian-preferred time (ECs: mean±*SEM*_*morning*_, 1.84±0.26µV, mean±*SEM*_*evening*_, 1.46±0.25µV; LCs: mean±*SEM*_*morning*_, 1.86±0.28µV, mean±*SEM*_*evening*_, 2.54±0.42µV) (Fig. 4c,d). We found a similar trend of P300 amplitude at the P3 electrode (Supplementary Fig. S3). Together, these results highlight the relationship between the circadian-preferred time and recruitment of motor learning-specific electrophysiological processes that are associated with performance. It should be noted thought that ERP amplitude enhancements at the circadian-preferred time could reflect enhanced learning, but also the faster frequency of movements caused by learning-related faster RT.

We also explored the association between motor learning, and plasticity by calculating correlations between MEPs amplitudes and motor sequence learning. Briefly, we found a positive correlation between anodal tDCS effects (MEP amplitude enhancement) and sequence learning (block 6 - 5 RT difference) in the evening for LCs (*r*=0.543, *p*=0.017). No significant correlation between sequence learning and tDCS-induced plasticity was found in ECs (for detailed results see Supplementary Material).

### Impact of chronotype on behavioural and electrophysiological correlates of cognition

Our results also indicated chronotype-specific variations of cortical excitability. Due to the links between cortical excitability, especially glutamate and GABA regulating, and cognitive processes^13, 47^, we next sought to determine the impact of chronotype on higher cognitive functions (e.g. working memory, attention), and respective ERP components as physiological indicators of information processing. All participants conducted the 3-back letter (working memory task), Stroop and AX-Continuous Performance tasks (AX-CPT) (attention tasks), during EEG recording at their circadian-preferred and non-preferred times (see Methods). Age, gender, and BMI did not covariate with the task outcome measures in the ANOVA analyses (Table 3).

For working memory performance, the ANOVA results revealed a significant chronotype**×**daytime interaction (*F*_1_ =10.34, *p*=0.003; *η*p^2^ =0.27) for the N-back hits, which is the primary outcome of interest in this task (Table 3). Post *hoc* Student’s t-tests showed a significantly enhanced WM performance in the morning for ECs and evening for LCs (Fig. 5a). The percentage of hits for ECs in the morning and evening was 65.39% and 57.15% respectively and for LCs in the evening and morning was 72.71% and 62.65 respectively. The same response pattern was observed for the electrophysiological correlates of N-back task performance. Additionally, we calculated the sensitivity index *d* (or d prime) which represents the proportion of hits rate minus the proportion of false alarm rate. A significant interaction of chronotype**×**daytime (*F*_1_ =10.82, *p*=0.003; *η*p^2^ =0.28) was found with no main effect of chronotype or time of day (Table 2). Post *hoc* Student’s t-tests showed a significantly enhanced *d* prime index in the morning for ECs and evening for LCs (Fig. 5a). Next, we found a significant interaction of chronotype**×**daytime in the P300 component at electrode Pz (*F*_1_ =12.39, *p*<0.001; *η*p^2^ =0.31), and Cz (*F*_1_ =11.07, *p*=0.002; *η*p^2^ =0.29) which is an indicator of memory-updating processes^48^. Post *hoc* comparisons via Student’s t-tests showed that performance during the circadian-preferred time was related to a larger P300 amplitude under the Pz electrode in both groups (ECs: mean±*SEM*_*morning*_, 3.37±0.50µV, mean±*SEM*_*evening*_, 2.43±0.47µV; LCs: mean±*SEM*_*morning*_, 1.76±0.48µV, mean±*SEM*_*evening*_, 2.75±0.39µV) (Fig. 5c). The same trend was observed for electrode Cz (Supplementary Fig. S4a,b).

**Fig. 5.**
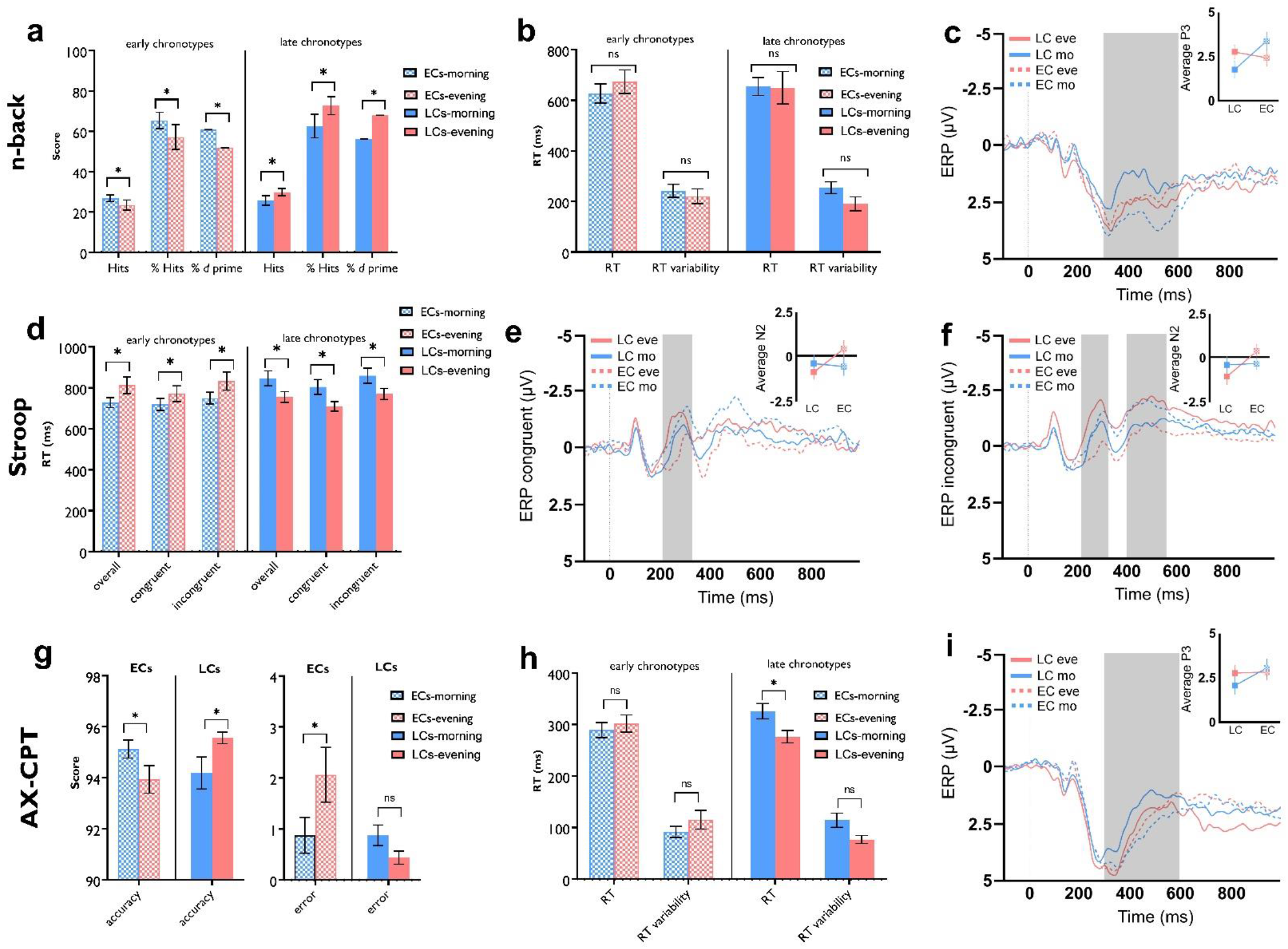
The impact of chronotype on working memory and attention. **a**, ECs had more correct responses and committed fewer errors in the morning vs evening (*t*=2.53, *p*=0.023), whereas LCs showed the reverse pattern of results (*t*=2.36, *p*=0.032) in the 3-back working memory task. Both groups showed the same pattern for the percentage of working memory accuracy and *d* prime (*t*_*ECs*_ =2.55, *p*=0.022; *t*_*LCs*_ =2.49, *p*=0.025) showing a significantly higher accuracy percentage and d prime value at their circadian-preferred time **b**, RT was not significantly different across time of day in the groups (*t*_*ECs*_ =1.13, *p*=0.275; *t*_*LCs*_ =0.18, *p*=0.858). **c**, In line with the behavioural results, ECs displayed a larger P300 in the morning compared to the evening (*t*=3.62, *p*=0.003) and LCs showed a larger P300 in the evening (*t*=2.27, *p*=0.038) at electrode Pz. **d**, Both, ECs (*t*_*morning*_ =1.91, *p*=0.074; *t*_*evening*_ =5.88, *p*<0.001) and LCs (*t*_*morning*_ =4.32, *p*=0.001; *t*_*evening*_ =5.60, *p*<0.001) displayed a stronger Stroop interference effect (RT_incongruent_ -RT_congruent_) at their circadian *non-preferred* time. RT of the congruent, incongruent and overall trials were significantly slower again at the circadian *non-preferred* time (ECs: *t*_*con*_ =1.66, *p*=0.117; *t*_*incon*_ =2.63, *p*=0.019; *t*_*overall*_ =3.43, *p*=0.004; LCs: *t*_*con*_ =4.13, *p*=0.001; *t*_*incon*_ =5.26, *p*<0.001; *t*_*overall*_ =5.17, *p*<0.001). **e, f**, The N200 amplitude at electrode Fz was calculated in the Stroop stage for both, congruent and incongruent trials. It was larger for both groups at their circadian-preferred times, but the difference was significant only in ECs (*t*_*con*_ =4.81, *p*=0.001; *t*_*incon*_ =2.65, *p*=0.018). However, chorotypes had a significant effect on the “morning vs evening” N200 amplitude difference (see results). For the N450 component details see Supplementary Fig. 4S,f. **g, h**, In the AX-CPT, both groups showed enhanced sustained attention, as indexed by higher accuracy (ECs: *t*=2.64, *p*=0.018; LCs: *t*=2.62, *p*=0.019) at their circadian preferred times. The respective RT difference was significant in LCs only (*t*=4.22, *p*=0.001). **i**, The P300 difference between morning and evening was significantly larger only for LCs (*t*=2.63, *p*=0.019). All pairwise comparisons are calculated using *Student’s* t-test. *n*=32 (16 per group). All error bars represent the s.e.m. across participants. eve = evening; mo = morning; P3 = P300 component; N2 = N200. Asterisks [*] represent statistically significant differences.

For Stroop task performance, a similar trend of response was observed. We found a significant chronotype**×**daytime interaction on overall RT (*F*_1_ =22.37, *p*<0.001; *η*p^2^ =0.45), RT of congruent trials (*F*_1_ =15.70, *p*<0.001; *η*p^2^ =0.36) and RT of incongruent trials (*F*_1_ =24.62, *p*<0.001; *η*p^2^ =0.47). The main effects of time of day and chronotype were significant neither (Fig. 5, Table 3). Post *hoc* comparisons of RTs revealed a significantly less Stroop effect (faster RT to incongruent trials) at the circadian-preferred time in both groups (Fig. 5d). This indicates less Stroop interference in participants when they were conducting the task at their circadian-preferred time. The same pattern of results was observed for RT variability in the Stroop block. The results of the ANOVA showed a significant interaction of chronotype**×**daytime (*F*_1_ =22.47, *p*<0.001; *η*p^2^ =0.45) and post *hoc* comparisons revealed a significantly reduced variability of RT in the Stroop block at the circadian-preferred time for both groups (ECs: *t*=2.81, *p*=0.013; LCs: *t*=4.62, *p*<0.001). Performance accuracy was not significantly affected by chronotype. The N200 and N450 are two prominent ERP markers related to Stroop conflict and are observed at frontocentral and centroparietal regions^49, 50^. The less Stroop effect we observed at the circadian-preferred times was associated with higher N200 and N450 amplitudes which are indicative of higher selective attention and discriminating conflicting stimuli. The results of ANOVA for the N200 amplitudes showed a significant interaction of chronotype**×**daytime on overall (*F*_1_ =22.47, *p*<0.001; *η*p^2^ =0.45), congruent (*F*_1_ =17.12, *p*<0.001; *η*p^2^ =0.39) and incongruent (*F*_1_ =7.17, *p*=0.012; *η*p^2^ =0.21) trials for the electrodes Fz. The N200 component of both congruent and incongruent trials over the Fz electrode was larger at the circadian preferred times in both groups (Fig. 5e,f), but the respective difference was significant only in ECs (incongruent: mean±*SEM*_*morning*_, -0.38±0.31µV, mean±*SEM*_*evening*,_ 0.33±0.36µV; congruent: mean±*SEM*_*morning*_, -0.59±0.29µV, mean±*SEM*_*evening*,_ 0.41±0.38µV). However, when we compared the amplitude difference values from morning to evening, chronotype had a significant effect (incongruent: *F*_1_ =7.57, *p*=0.010; *η*p^2^ =0.21; congruent: *F*_1_ =14.49, *p*<0.001; *η*p^2^ =0.33) yielding higher negativity of N200 at circadian-preferred times in both groups. The same pattern of response was found for the Cz electrode and the N450 component (Supplementary Fig. S4c,d,e,f).

The next performed task was the AX-CPT which is a lower cognitive-demanding task for measuring sustained attention (see Methods). Analysis of the behavioural data showed a significant interaction of chronotype**×**daytime on both accuracy (*F*_1_ =14.16, *p*<0.001; *η*p^2^ =0.34) and RT (*F*_1_ =19.39, *p*<0.001; *η*p^2^ =0.41). Both groups performed more accurately when the task was conducted at their circadian-preferred time. With regard to RT, only the LCs showed a significantly faster RT at their circadian-preferred time (Fig. 5g,h). The P300 serves as an attentional index to stimulus and memory storage, which are required in the AX-CPT. Analysis of this ERP component showed a significant interaction of chronotype**×**daytime in the P300 component at electrode Pz (*F*_1_ =5.33, *p*=0.028; *η*p^2^ =0.16). Post *hoc Student’s* t-tests indicated that the circadian-preferred time was significantly related to a larger P300 amplitude only in the LCs (LCs: mean±*SEM*_*morning*_, 2.04±0.49µV, mean±*SEM*_*evening*_, 2.74±0.42µV; ECs: mean±*SEM*_*morning*_, 3.03±0.50µV, mean±*SEM*_*evening*_, 2.80±0.44µV) (Fig. 5i).

### No difference in sleepiness rating and physiological marker of sleep pressure across groups

All participants had moderate chronotypes which reduces the variability of sleep-wake cycle, however, this can interfere with chronotype-specific effects on brain physiology and cognition in case of higher sleep pressure in either group at non-preferred times. To resolve this, we controlled the sleep-wake cycle and sleep duration of both groups within a defined range (see methods for details) and rated participants’ sleepiness with the Karolinska Sleepiness Scale before each test session. All participants had at least 8 h of sleep before each session (see methods). The results of ANOVA showed no significant interaction of chronotype**×**daytime (*F*_1_ =1.03, *p*=0.325) or main effects of chronotype and time of day. This indicates that there was no significant difference between the sleepiness ratings in or between both groups across different times of day. Furthermore, we analyzed resting EEG theta oscillations, which is an objective marker of sleep pressure^51^ in both groups in the morning and evening. There were no significant differences in the theta oscillations at frontocentral electrodes when we compared both groups in the morning (*t*_*Cz*_ =0.48, *p*>0.999) and evening (*t*_*Cz*_ =1.80, *p*=0.433) (For Fz and Pz see Supplementary material). No significant differences were neither observed in each group across different times of the day. The same pattern of results was observed for alpha oscillations (Supplementary material).

## Discussion

Previous studies have shown that chronotype has distinctive behavioural, physical, and genetic manifestations in humans^2^. Here we showed a converging impact of chronotypes on time-of-day dependent behavioural/cognitive performance of healthy individuals, and demonstrated the physiological foundations of these effects by daytime-dependent cortical excitability, neuroplasticity, and brain information processing processes (Fig. 6).

**Fig. 6.**
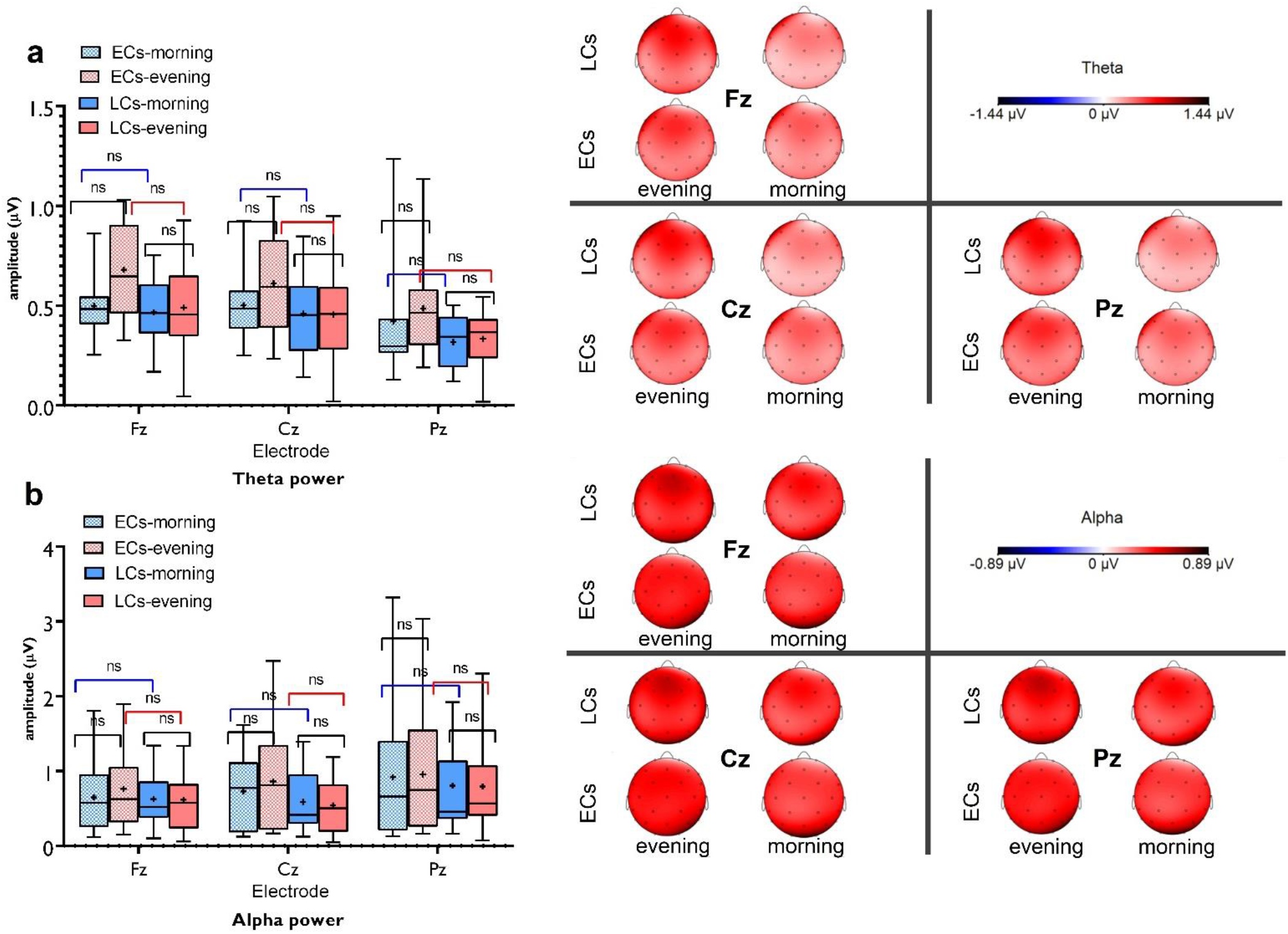
Resting-EEG theta and alpha oscillations at circadian-preferred and non-preferred time for ECs and LCs. (**a**) The results of 3 (Fz, Cz, Pz electrodes) × 2 (chronotype) × 2 (daytime) ANOVA showed no significant interaction of electrode×chronotype×daytime (*F*_1.36_ =1.65, *p*=0.207) or electrode×chronotype (*F*_1.90_ =0.20, *p*=0.806) or electrode×daytime (*F*_1.36_ =2.17, *p*=0.142) on theta oscillations. The interaction of chronotype×daytime was marginally significant (*F*_1_ =4.65, *p*=0.040). Post hoc comparisons showed no significant difference of theta oscillations between groups in the morning (*t*_Fz_ =0.36, *p*>0.999; *t*_Cz_ =0.53, *p*>0.999; *t*_Pz_ =1.22, *p*>0.999) and evening (*t*_Fz_ =2.42, *p*=0.097; *t*_Cz_ =1.88, *p*=0.371; *t*_Pz_ =1.86, *p*=0.382). (**b**) For alpha oscillations, no significant interaction of electrode×chronotype×daytime (*F*_1.14_ =0.30, *p*=0.627) or electrode×chronotype (*F*_1.36_ =1.17, *p*=0.302), electrode×daytime (*F*_1.14_ =0.35, *p*=0.615) or chronotype×daytime (*F*_1_ =1.68, *p*=0.204) were found. Pairwise comparisons are calculated by post hoc t-tests (paired, two-sided, *p*<0.05). *n*=30 (15 ECs, 15 LCs). Data are presented as mean values±SEM. The horizontal bar shows the median, the + shows the mean, the upper and lower boundaries show the 25th and 75th percentiles, respectively and the whiskers show the 5-95 percentile. ECs = early chronotypes; LCs = late chronotypes; ns = nonsignificant; µV= microvolt. [*] indicates a significant difference.

**Fig. 7.**
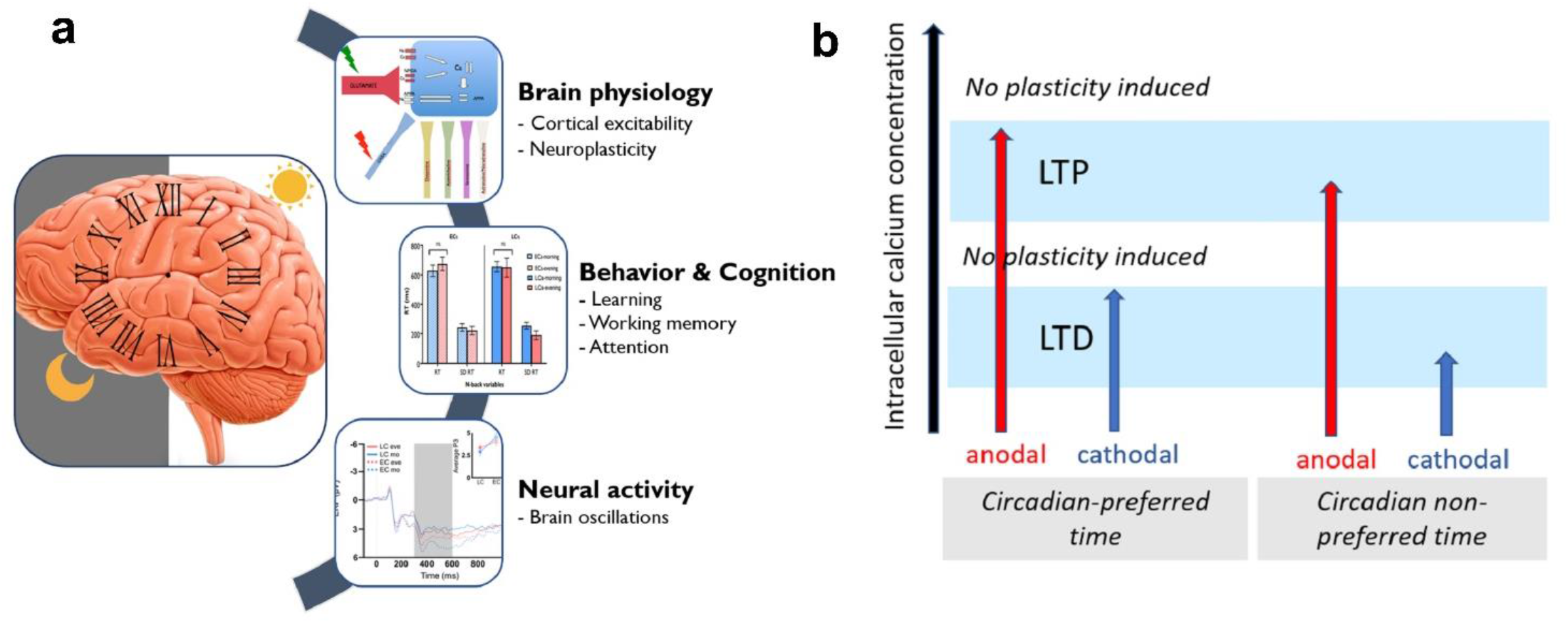
**a**, A schematic illustration depicting the converging impact of chronotype on brain physiology, behavior, and cognition. **b**, Proposed mechanism of the neuroplasticity induction at circadian-preferred and non-preferred time based on the association between intracellular calcium concentration (x-axis) and induction of tDCS-induced neuroplastic changes. Gradual enhancement of calcium concentration can either induce LTD, LTP or no plasticity. The LTP-like plasticity induced by tDCS is linked to higher intracellular calcium concentration while LTD-like plasticity induced by tDCS is linked to a lower intracellular calcium concentration. It can be assumed that intracellular calcium concentration at the optimal time of the day is at an optimal level leading to stronger tDCS-induced LTP/D like plasticity. This would be at least partially related to the higher glutamatergic and lower GABAergic activation at the circadian preferred time, shown by cortical excitability data.

The significantly higher cortical facilitation and lower cortical inhibition at the circadian-*preferred* time in both chronotypes argues for specific differences of cortical physiology mediated by chronotype and time of day. Specifically, our results suggest that at the circadian-preferred time, intracortical facilitation is enhanced predominantly by increased activity of glutamatergic synapses. Conversely, cortical inhibition was significantly pronounced at the circadian *non*-preferred time presumably through enhanced GABAergic activation. These results are in accordance with evidence from animal studies, which show a circadian regulation of GABA and glutamate^52-54^ highlighting the importance of excitatory and inhibitory systems in cortical excitability. Specifically, GABA is an important synchronizer of the suprachiasmatic nucleus, the major structure involved in circadian rhythms, whose activity varies throughout the day^55^. Studies in humans identified regulation of the GABAergic system, and respective alterations of cortical inhibition/facilitation within the circadian cycle as well^5, 28^. Yet, *chronotype* effects on brain physiology were not specifically addressed in previous studies. Here we demonstrated *chronotype*-specific modulation of cortical excitability for the first time.

In line with these results, we found that LTP/LTD-like neuroplasticity, which depends on the glutamatergic and GABAergic systems, is modulated by chronotype too. Evidence from primary motor cortex models in humans and animals show that tDCS-induced plasticity is driven by activation of NMDA receptors, and gated by reduction of GABA activity^20^. Specifically, anodal stimulation LTP-like after-effect is thought to be caused by a major enhancement of NMDA receptor activity, while cathodal stimulation-generated LTD-like plasticity is suggested to involve a minor enhancement of NMDA receptors, driven by reduction of glutamate. Moreover, both kinds of plasticity seem to require reduction of GABA activity^23, 41^. We argue that chronotype-specific differences of glutamatergic facilitation and GABAergic inhibition at circadian-preferred and non-preferred times, as described above, can explain the plasticity differences we obtained. A brain state of enhanced glutamatergic activity, and reduced GABAergic inhibition, as present in the morning hours for ECs, and the evening hours in LCs, would facilitate plasticity induction presumably via the optimal intracellular calcium concentration which determines the plasticity zones (Fig. 6b). In line with this, we demonstrated for the first time that active tDCS, compared to sham stimulation, induced LTP/LTD-like neuroplasticity depending on the circadian preference, which was correlated with cortical excitability measures (Supplementary material). These findings are consistent with previous animal studies that revealed a strong effect of the circadian clock on hippocampal plasticity^56^ and complement those of human studies that showed that plasticity response to a given paired-associative stimulation is regulated by circadian rhythms^16^.

The important question here is whether these chronotype-specific differences in brain plasticity influence learning and memory formation that depend on brain plasticity as well^57^. Providing proof of this, we found that motor sequence learning and retention, and respective ERP components, follow the same chronotype-facilitating effect and plasticity induction and motor learning at circadian-preferred times were associated. This observation makes sense because tDCS-induced neuroplasticity in the motor cortex and behavioural motor learning share intracortical mechanisms^17^. Furthermore, evidence from magnetic resonance spectroscopy shows a learning-specific reduction of GABA concentration in the motor cortex during motor learning^42^. In agreement with this, we found less GABAergic cortical inhibition at the circadian-preferred time which was associated with stronger motor learning at the behavioural level. Together, these convergent findings establish an important impact of chronotype on motor cortex functionality from basic physiological functions (cortical excitability, LTP-like neuroplasticity) to behavioural and electrophysiological levels.

Cortical excitability alterations are expected to be associated with changes in behavioural and cognitive performance^13^. We found an impact of time-of-day on basic cognitive processing, including working memory and attentional functioning, indicating that chronotype-specific effects are extended to non-motor areas. Animal studies have demonstrated common mechanisms (e.g. neurotransmitter release, synaptic excitability, and neuronal activity) underlying circadian rhythm and memory formation^26^. For working memory, several circadian-related molecular mechanisms are proposed, including circadian clock-gated changes in glutamate receptor activity^58^. These mechanisms are in line with the enhanced cortical excitability caused by increased glutamatergic facilitation and LTP-like plasticity at the circadian-preferred time which we found in our excitability measures. Regarding attentional functioning, the peaks and troughs in circadian rhythms can differentially impact on performance^6^. Our data are in agreement with these findings by showing an association between physiological plasticity and excitability data, which are indicative of daytime- and chronotype-dependent differences of glutamate and GABA activity, and cognitive performance of chronotype at respective times of day (for details of respective correlations see Supplementary material). Moreover, task-specific ERP components were correlated with cognitive performance as well. The N200 and P300 components reflect stimulus identification/distinction and memory-updating processes,^48^ suggesting a link between circadian preference and the physiological foundation of cognitive processing. These effects were more clear for the tasks with a higher cognitive load, such as the 3-back letter task, in line with findings of cognitive load-dependency of circadian effects^6^.

The findings of this study have specific implications for the field of human neurophysiology and cognitive neuroscience as well as broad implications for human behavior in healthy and clinical populations. “Time-of-day” and chronotype are important, but less-studied determinants of cortical plasticity induction by NIBS techniques^13, 16^. Our results show that cortical excitability and neuroplasticity are chronotype-dependent in humans. It is tempting to speculate that chronotype could account for variability in the efficacy of NIBS, and might explain partially heterogeneous effects in previous studies. This assumption might be likewise relevant for the performance of various cognitive tasks and suggest screening or considering its modulatory effects. Being studied at the optimal time of day, having sufficient sleep and control of interfering factors might enhance homogeneity of results, as our data suggest. Beyond genetic determinants^2^, chronotype is dependent on social pressures and our modern lifestyle that is increasingly deviating from the 24-hr cycle. It has a clear effect on sleep timing as well^2^ which indicates its relevance for working and educational environments. Working at circadian-antagonistic time can disrupt the circadian cycle and thereby the shift workers’ health^59^. Learning materials and studying, which are dependent on learning, memory, and attention, can be hindered at circadian non-preferred times. At the clinical level, it can affect therapeutic efficacy of NIBS and other interventions. For example, the ability to learn novel motor skills is central for the rehabilitation after a stroke, which could vary depending on patients’ chronotype, and timing of rehabilitation. Furthermore, circadian disruption is linked to cardiometabolic disorders and the pathophysiology of neurodegenerative diseases (e.g. Parkinson’s disease, Alzheimer’s disease)^59^ and should be considered in personalized and precision medicine, and timing of interventions for higher efficacy.

It is important to consider that our measurement times were fixed across groups and were not individually adapted to each subject. The main reason lies in our study question which was whether brain physiological parameters and cognitive performance differ at different times of day due to *chronotype* but not different levels of sleep pressure observed in different chronotypes. Therefore, we had to pick a fixed time and at the same time assure that there is no sleep pressure for each group at their nonpreferred time of day. To this end, we controlled timing to go to bed and sleep duration before each experimental session and measures potential sleep pressure by both subjective sleepiness scale and objective EEG oscillations as a marker of sleep pressure^51^. The non-significant difference between the sleepiness ratings before each session across groups, and more importantly data of theta and alpha oscillations at the experimental time indicates that potential sleep loss or different sleep-wake cycles do not account for the observed effects. Our measurement times were selected based on the “chronotype-based paradigm”^6^ and sleep-wake cycle and level of activity in early and late chronotypes^60^.

Our physiological measures were based on the motor system and indirect measures of the involved neurotransmitters, yet they suggest systematic effects of chronotype at the whole-brain level. Interestingly, chronotype-specific individual differences in brain anatomy (e.g. grey matter volume) have been described by some studies^61^, which could affect tDCS-induced neuroplasticity induction either in a facilitatory or inhibitory way. It is unlikely that differences of brain structure were the driving force of the results because both groups showed enhanced plasticity at specific times of the day congruent with their respective chronotype. Furthermore, although there was no significant difference between baseline MEPs and % of MSO across conditions in both groups, which supports the reliability of the acquired data, the use of neuronavigation for stimulation of the motor cortex might have been advantageous to further enhance reproducibility of the TMS coil position. Finally, although NIBS allows us to causally modify and induce neuroplasticity in humans non-invasively, it is worth acknowledging that the evidence for synaptic plasticity induction by NIBS, including tDCS, comes from animal and pharmacological studies, and is thus indirect^62-64^. In conclusion, our results show a relevant impact of circadian preference on learning and cognition including memory formation and attentional functions as well as brain physiology underlying these cognitive processes including cortical excitability, neuroplasticity, and electrophysiological correlates of cognitive processes.

## Acknowledgments

We greatly appreciate the comments and support offered by Prof. Barbara Griefahn. MAN is supported by a grant from the German Ministry of Research and Education (GCBS grant 01EE1403C). We appreciate Dr. Fatemeh Yavari, Dr. Asif Jamil, and Mr. Tobias Blanke, for their technical support.

## Author contributions

**M.A.S**. conducted the experiments, collected and analyzed the data, wrote the paper and designed illustrations. **M.W**. and **M.A.S**. analyzed EEG data analysis and illustrated EEG illustrations.

**M.M.S**. and **E.GH**. conducted 3D computational modeling, designed and illustrated figures. **M-F.K**. and **M.A.N**. conceived and designed the experiments and revised the final draft. All authors contributed to the final drafting of the work.

